# Neutrophil elastase inhibitor sivelestat ameliorates gefitinib-naphthalene-induced acute pneumonitis in mice

**DOI:** 10.1101/108464

**Authors:** Hironori Mikumo, Toyoshi Yanagihara, Naoki Hamada, Eiji Harada, Saiko Ogata-Suetsugu, Chika Ikeda-Harada, Masako Arimura-Omori, Kunihiro Suziki, Tetsuya Yokoyama, Yoichi Nakanishi

## Abstract

**Background and objective:** Gefitinib, an epidermal growth factor receptor-tyrosine kinase inhibitor (EGFR-TKI), is an effective therapeutic agent for non-small cell lung cancer with EGFR mutations. It can cause severe acute pneumonitis in some patients. We previously demonstrated that mice with naphthalene-induced airway epithelial injury developed severe gefitinib-induced pneumonitis and that neutrophils played important roles in the development of the disease. This study aimed to investigate the effects of the neutrophil elastase inhibitor sivelestat on gefitinib-induced pneumonitis in mice.

**Methods:** C57BL/6J mice received naphthalene (200 mg/kg) intraperitoneally on day 0. Gefitinib (250 or 300 mg/kg) was orally administered to mice from day −1 until day 13. Sivelestat (150 mg/kg) was administered intraperitoneally from day 1 until day 13. Bronchoalveolar lavage fluid (BALF) and lung tissues were sampled on day 14.

**Results:** Sivelestat treatment significantly reduced the protein level, neutrophil count, neutrophil elastase activity in BALF, and severity of histopathologic findings on day 14 for mice administered with 250 mg/kg of gefitinib. Moreover, sivelestat treatment significantly improved the survival of mice administered with 300 mg/kg of gefitinib. Conclusions: These results indicate that sivelestat is a promising therapeutic agent for severe acute pneumonitis caused by gefitinib.

**Summary statement:** Neutrophil elastase inhibitor sivelestat is a promising therapeutic agent for severe acute pneumonitis caused by gefitinib.

## INTRODUCTION

Gefitinib, an epidermal growth factor receptor-tyrosine kinase inhibitor (EGFR-TKI), is an effective therapeutic agent for non-small cell lung cancer with EGFR mutations (Harada et al., 2011; Mok et al., 2009). It can cause severe acute pneumonitis in some patients. Characteristics of patients who developed interstitial pneumonia included old age, poor performance status, a history of smoking, and preexisting interstitial pneumonia (Harada et al., 2011; Kudoh et al., 2008).

Injuries to respiratory epithelium and alveolar epithelial cells are regarded as the initial phenomena of various respiratory illnesses, such as acute respiratory distress syndrome, interstitial pneumonia, and chronic obstructive pulmonary disease.

Neutrophil elastase is a protease produced by neutrophils. Excessive neutrophil elastase can cause lung tissue damage by direct cytotoxicity to endothelial and epithelial cells and by degradation of key structural elements of connective tissue, such as elastin, collagen, and proteoglycan (Yamada et al., 2011; Lee and Downey, 2001).

Sivelestat, a small molecule (529 Da), is a neutrophil elastase inhibitor developed and produced by Ono Pharmaceutical Company in Japan (Aikawa et al., 2011). In the animal models of acute lung injury (ALI), the beneficial effects of sivelestat have been reported in bleomycin-induced inflammation and streptococcus pneumonia (Yuan et al., 2014; Yamada et al., 2011). A phase 3 study in Japan demonstrated that sivelestat improved the investigator assessment of pulmonary function and significantly reduced the duration of intensive care required for patients with ALI associated with systemic inflammatory response syndrome (SIRS) (Tamakuma et al., 2004). In 2002, sivelestat was approved in Japan for the treatment of ALI associated with SIRS (Aikawa et al., 2011). After approval, a phase 4 study indicated that it contributed to early weaning from mechanical ventilation (Aikawa et al., 2011). The beneficial effects of sivelestat have also been reported in several other models, including lipopolysaccharide (LPS)-induced lung inflammation, ozone-induced airway response, and bleomycin-induced pulmonary fibrosis (Yuan et al., 2014; Matsumoto et al., 1999; Takemasa et al., 2012).

Pulmonary stem cells are important for tissue recovery. The club cell is a type of pulmonary stem cell found in the distal airway. Stem cell abnormality is considered to promote chronic lung injury and lung fibrosis (Harada et al., 2011; Gazdhar et al., 2007). Naphthalene has club cell-selective cytotoxicity (Harada et al., 2011; Van Winkle et al., 1995; Stripp et al., 1995). Therefore, we previously used a naphthalene-induced lung injury model as an animal model containing a risk factor of gefitinib-induced pneumonia. We found that gefitinib administration after naphthalene treatment prolonged ALI with neutrophil infiltration on day 14. Laser capture microdissection and microarray analysis of the terminal bronchial epithelial cells showed upregulation of the genes included—*S100a8*, *S100a6*, *Stfa3*, *Trim23*, and *Mug1*, which are known to participate in inflammatory cell chemotaxis, activation, and migration (Harada et al., 2011; Raquil et al., 2008; Ozato et al., 2008). Although the precise mechanisms involved remain unclear, gefitinib treatment prolonged lung inflammation by upregulating neutrophil chemoattractant genes from peripheral epithelial cells. Therefore, we hypothesize that sivelestat plays a protective role against gefitinib-induced lung injury by inhibiting neutrophilic inflammation.

## RESULTS

### Sivelestat improved the survival rate of gefitinib-induced pneumonitis in mice

The administration of 300-mg/kg gefitinib with 150-mg/kg sivelestat following naphthalene significantly improved the survival rate on day 14 compared with that of 300 mg/kg gefitinib following naphthalene (Figure 2).

**Figure 1.**
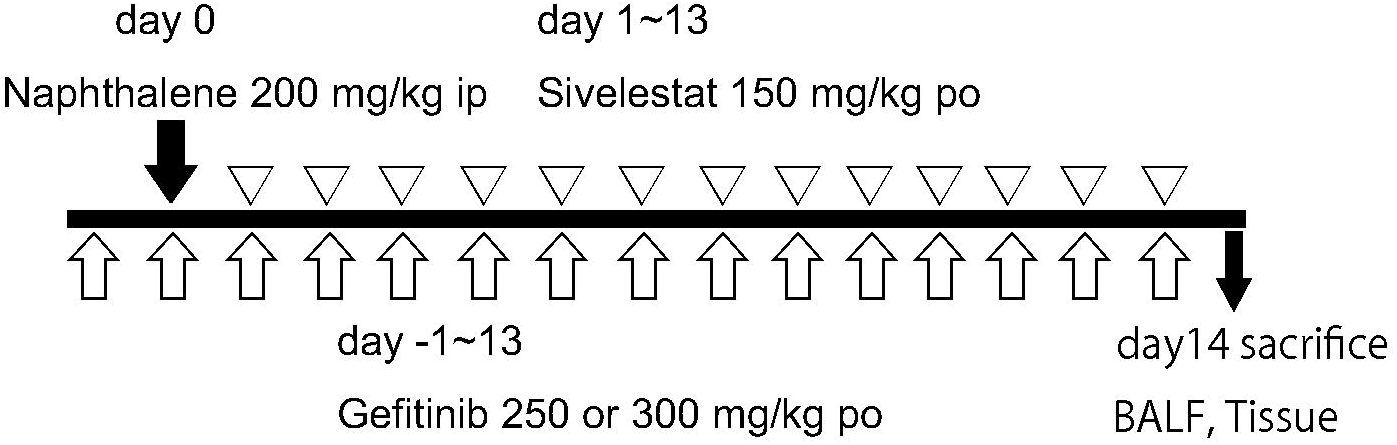
Experimental scheme detailing the administration of naphthalene, gefitinib, and sivelestat. po: oral administration, ip: intraperitoneal administration

**Figure 2.**
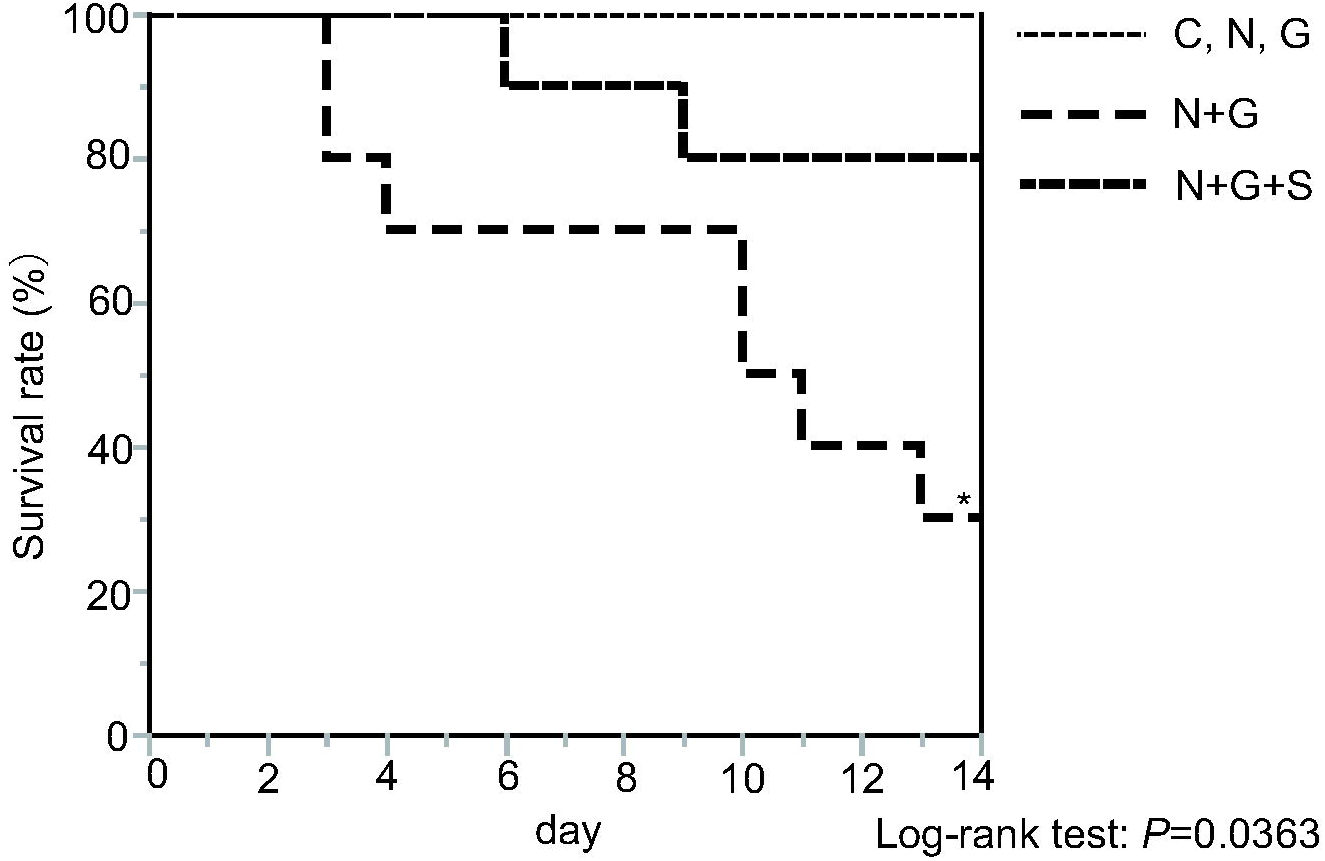
Kaplan–Meier survival curve Survival study of mice with the administration of 300 mg/kg gefitinib and 150 mg/kg sivelestat following naphthalene improved the survival rate compared with that of 300 mg/kg gefitinib following naphthalene (*n* = 10). \**P* < 0.05.

### Sivelestat ameliorated the loss of body weight of gefitinib-induced pneumonitis in mice

We measured mice body weight to determine the general influence of ALI. Weight loss is a good marker for the severity of naphthalene-induced lung injury (Harada et al., 2011; Verschoyle et al., 1997). On day 7, the body weights of mice treated with naphthalene alone were significantly decreased; however, by day 14, the body weights returned to the level of the control injected with corn oil. On day 14, body weights of mice treated with 250 mg/kg gefitinib following naphthalene remained significantly decreased compared with that of mice treated with naphthalene alone, whereas the body weights of mice treated with 250 mg/kg gefitinib and 150 mg/kg sivelestat following naphthalene were significantly increased compared with that of mice treated with 250 mg/kg gefitinib following naphthalene (Figure 3).

**Figure 3.**
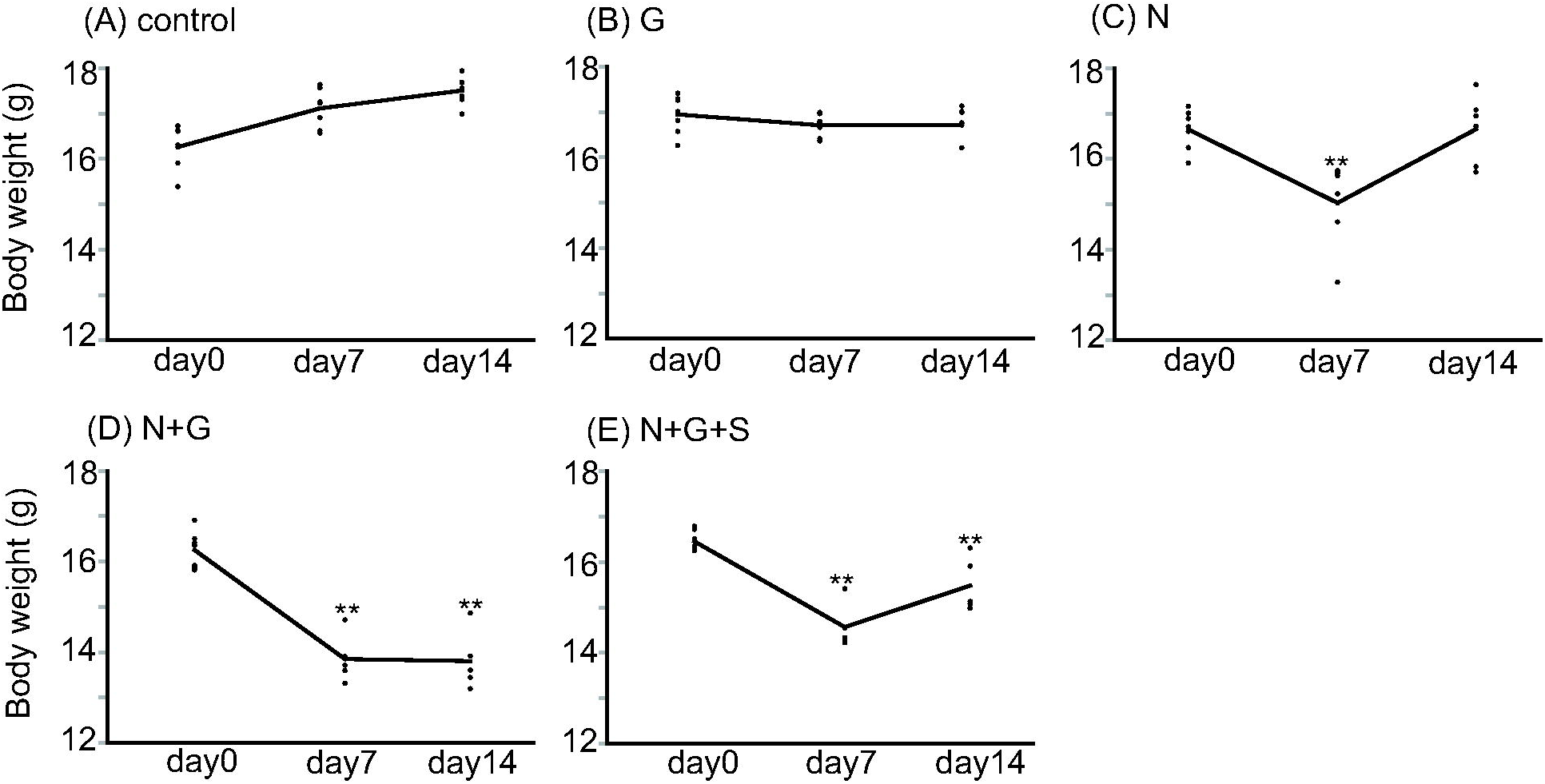
Changes in mice body weight over time. Body weights of mice treated with naphthalene alone were significantly decreased on day 7; however, by day 14, the body weights returned to the level of the control. On day 14, the body weights of mice treated with 250 mg/kg gefitinib following naphthalene remained significantly decreased compared with that of mice treated with naphthalene alone, whereas the body weights of mice treated with 250 mg/kg gefitinib and 150 mg/kg sivelestat following naphthalene were significantly increased compared with those of mice at treated with 250 mg/kg gefitinib following naphthalene. Compared with control group at the same point (*n* = 7). \**P* < 0.05, \*\**P* < 0.01.

### Sivelestat ameliorated the lung inflammation of gefitinib-induced pneumonitis in mice

#### Histopathological examination

We used histologic cell analysis to examine the degree of inflammatory cell infiltration. Naphthalene alone induced neutrophil infiltration on day 7 but not on day 14 in the lung tissue, as previously described (Harada et al., 2011). The administration of 250 mg/kg gefitinib following naphthalene aggravated neutrophil infiltration and induced alveolar hemorrhage on day 14 (Figure 4A). In contrast, the administration of 250 mg/kg gefitinib with 150 mg/kg sivelestat following naphthalene significantly decreased the pathologic grade compared with that of 250 mg/kg gefitinib following naphthalene (Figure 4B).

**Figure 4.**
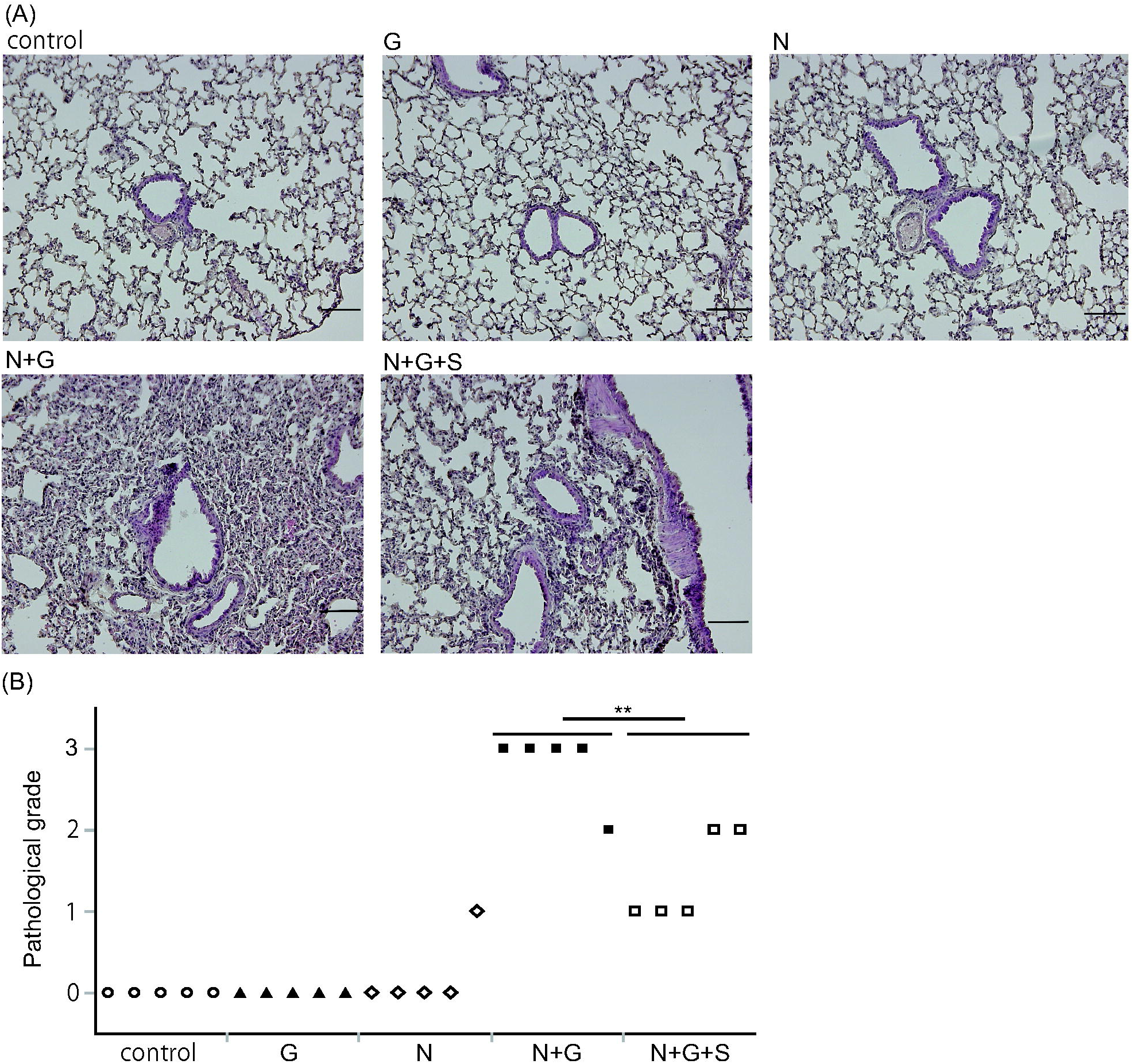
Histologic assessment of lung tissues on day 14. (A) Hematoxylin and eosin staining. The administration of 250 mg/kg gefitinib following naphthalene significantly induced neutrophil infiltration and acute lung injury. The administration of 250 mg/kg gefitinib and 150 mg/kg sivelestat following naphthalene improved neutrophil infiltration compared with that with 250 mg/kg gefitinib following naphthalene (*n* = 5), Scale bars: 100 μm. (B) Pathologic grade of lung tissues on day 14 (*n* = 5). \*\**P* < 0.01.

#### BALF analysis

On day 14, the number of neutrophils, total cell count, and protein concentration in BALF of mice treated with 250 mg/kg gefitinib following naphthalene were significantly increased compared with those of mice treated with naphthalene alone. On day 14, the administration of 250 mg/kg gefitinib and 150 mg/kg sivelestat following naphthalene significantly decreased the number of neutrophils, total cell count, and protein concentration in BALF compared with that with 250 mg/kg gefitinib following naphthalene (Figure 5).

**Figure 5.**
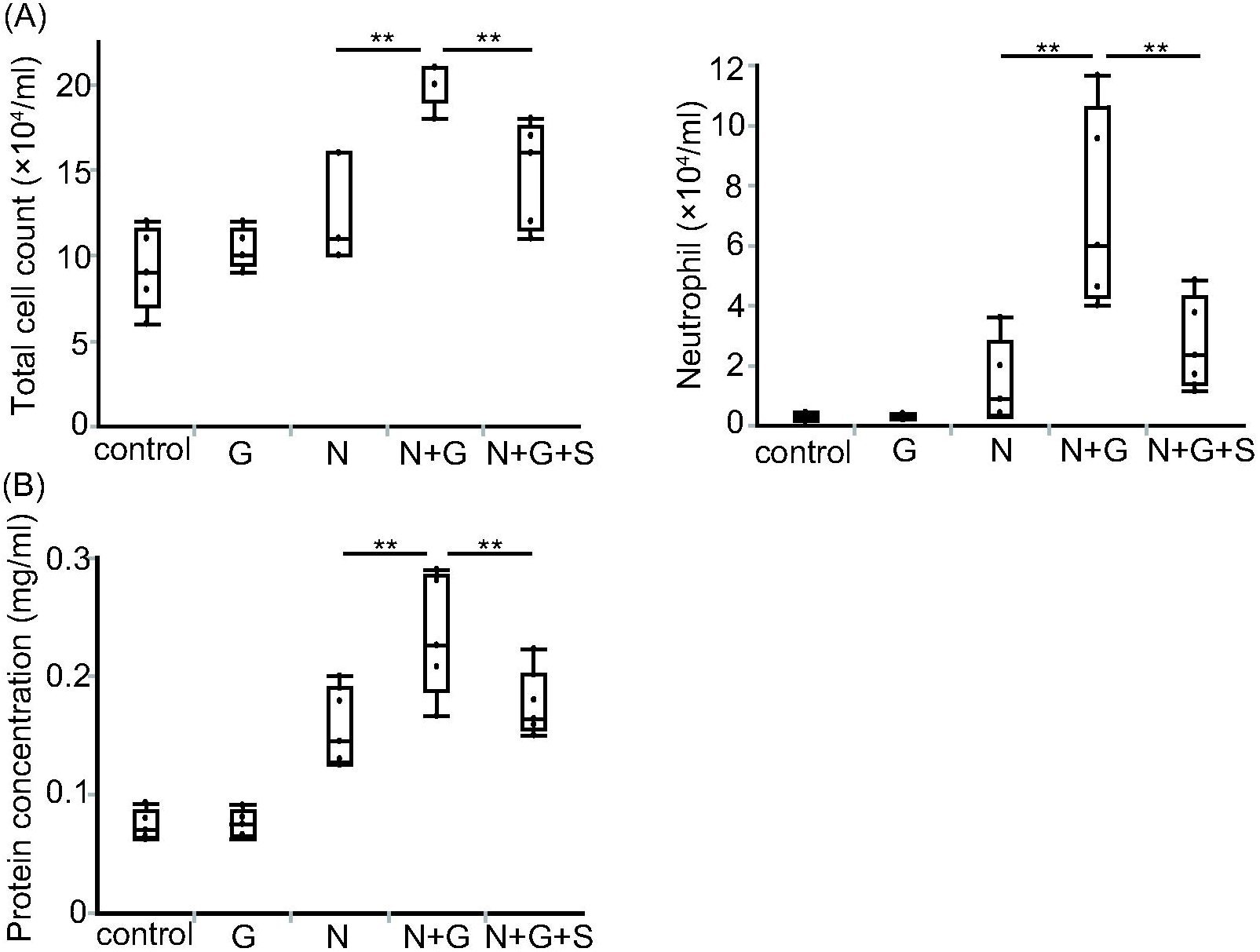
Bronchoalveolar lavage fluid (BALF) analysis on day 14 (A) The administration of 250 mg/kg gefitinib following naphthalene significantly induced the upregulation of total cell count and neutrophil recruitment, which were decreased by administration of 250 mg/kg gefitinib and 150 mg/kg sivelestat following naphthalene (*n* = 5). \*\**P* < 0.01. (B) The administration of 250 mg/kg gefitinib following naphthalene significantly induced the upregulation of protein concentration, which was decreased by the administration of 250 mg/kg gefitinib and 150 mg/kg sivelestat following naphthalene (*n* = 5). \*\**P* < 0.01.

In addition, the administration of 250 mg/kg gefitinib and 150 mg/kg sivelestat following naphthalene significantly decreased the level of IL-8 (Figure 6A) and neutrophil elastase activity (Figure 6B) in BALF compared with those resulting from the administration of 250 mg/kg gefitinib following naphthalene.

**Figure 6.**
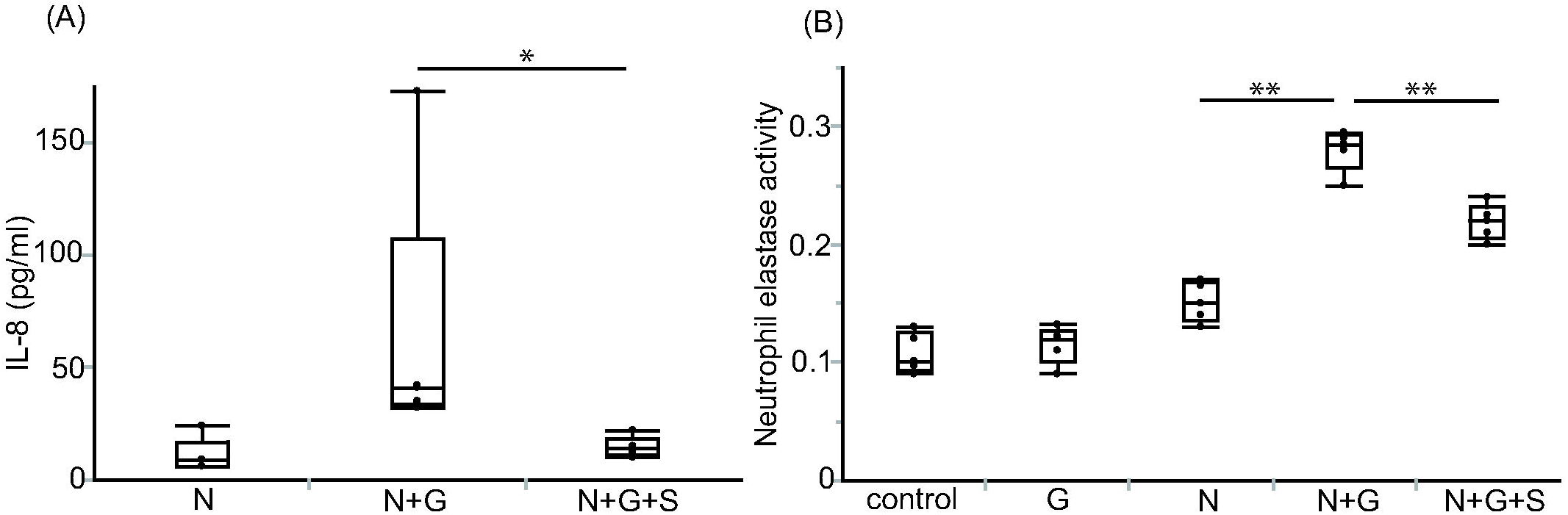
Interleukin-8 (IL-8) and neutrophil elastase activity in bronchoalveolar lavage fluid (BALF) on day 14 (A) IL-8 was increased by the administration of 250 mg/kg gefitinib following naphthalene and was decreased by administration of 250 mg/kg gefitinib and 150 mg/kg sivelestat following naphthalene (*n* = 5). \**P* < 0.05. (B) Neutrophil elastase was increased by the administration of 250 mg/kg gefitinib following naphthalene and was decreased by administration of 250 mg/kg gefitinib and 150 mg/kg sivelestat following naphthalene (*n* = 5). \*\**P* < 0.01.

## DISCUSSION

As we reported previously, the presumed sequence of the mechanism of gefitinib– naphthalene pneumonitis is as follows: 1) the upregulation of neutrophil chemoattractant genes in bronchiolar epithelial cells; 2) neutrophil migration into alveolar space and interstitial tissues, and; 3) release of neutrophil elastase from neutrophils, resulting in lung tissue damage.

IL-8 is produced by alveolar epithelial cell line (A549), airway epithelial cells, and inflammatory cells, such as macrophages and neutrophils. IL-8 is a well-known neutrophilic chemoattractant. Some studies have reported that gefitinib induces the production of IL-8 from alveolar epithelial cell line (A549). Neutrophil elastase induces the release of IL-8 from bronchial epithelial cells (Yamada et al., 2011; Nakamura et al., 1992), which in turn recruits additional neutrophils. Our study indicated that administration of sivelestat decreased neutrophil elastase, which in turn inhibited the level of IL-8 in gefitinib–naphthalene-induced pneumonitis. In other words, sivelestat treatment could halt the negative spiral of lung injury. A limitation of the current study was that we could not identify the IL-8 producing cells targeted by sivelestat; therefore, further studies are required.

A significant reduction in the number of club cells has been observed in the airway epithelium of chronic tobacco smokers (Nomori et al., 1994). The naphthalene-induced club cell injury in a mice model may be representative of patients at a high risk of gefitinib-induced pneumonitis. This suggests that the presence of peripheral airway damage may increase the susceptibility of patients with lung cancer to interstitial pneumonia during treatment with gefitinib.

In conclusion, we demonstrated a treatment strategy for ALI caused by gefitinib. Currently, new generation EGFR-TKIs, such as afatinib and osimertinib, are used for treating non-small cell lung cancer with EGFR mutations. It is well known that the new generation EGFR-TKIs also induce pneumonitis; therefore, the current study should be repeated for the new generation EFGR-TKIs.

## MATERIALS AND METHODS

### Animal treatment

The experiments were approved by the Committee on Ethics Regarding Animal Experiments of Kyushu University. C57BL/6 female mice (7 weeks old; SLC, Inc, Shizuoka Japan) were used in all experiments.

Naphthalene (Wako Pure Chemical Industries, Osaka, Japan) was injected intraperitoneally on day 0 (200 mg/kg). Gefitinib (Caymann Chemical, Arizona, USA) stirred into 1% Tween 80 (Wako) was daily administered orally on days −1 to 13. We administrated gefitinib at two doses: 1) 250 mg/kg as a tolerated dose and 2) 300 mg/day as the 50% lethal dose (LC_50_). The neutrophil elastase inhibitor sivelestat (Ono Pharmaceutical, Osaka, Japan) in saline was daily injected intraperitoneally on days 1 to 13 (150 mg/kg). A scheme of the administration schedule is shown in Figure 1.

### Histopathological evaluation

Histopathology was performed as previously described (Harada et al., 2011; Hamada et al., 2008).

The right lung was fixed in 10% buffered formalin and embedded in paraffin, and the lung sections were stained with hematoxylin and eosin. The pathological grade of inflammation in the whole area of the midsagittal was evaluated under ×200 magnification and determined according to the following criteria: 0 = no lung abnormality; 1 = presence of inflammation involving <25% of the lung parenchyma; 2 = lesions involving 25–50% of the lung; and 3 = lesions involving >50% of the lung.

### Bronchoalveolar lavage

The bronchoalveolar lavage (BAL) method and analysis was performed as previously described (Harada et al., 2011; Hamada et al., 2008). After counting the cell numbers in BAL fluid (BALF), cells were cytospun and stained with Diff-Quick for classification. The BALF supernatant was freeze-dried using a lyophilizer. The lyophilized samples were dissolved to determine total protein concentrations, interleukin-8 (IL-8), and neutrophil elastase activity. Total protein concentrations in BALF were measured using the Bio-Rad Protein Assay.

### ELISA for assessment of IL-8 in the BALF

The concentration of IL-8 in BALF was determined using mouse cytokine ELISA kits (R&D Systems, Minneapolis, MN, USA).

### Neutrophil elastase activity

Neutrophil elastase activity in BALF was determined using the highly neutrophil elastase-specific synthetic substrate *N*-methoxysuccinyl-Ala-Ala-Pro-Val *p*-nitroanilide. Briefly, samples were incubated in 0.1-M Tris-HCl buffer (pH 8.0) containing 0.5-M NaCl and 1-mM substrate for 24 h at 37 ºC. After incubation, *p*-nitroaniline was measured spectrophotometrically at 405 nm, considered to be a measure of neutrophil elastase activity (Hagio et al., 2004; Yanagihara et al., 2007).

### Statistical Analysis

The Student’s t-test was used for the comparison of body weight, number of BALF cells, protein concentration, IL-8 concentration, neutrophil elastase activity, histopathological grade, and survival curves. *P* < 0.05 was considered significant. Statistical analysis was performed in the statistical software package JMP version 11 (SAS Institute, Cary, NC).

## ACKNOWLEDGEMENTS

We thank S. Tamura for technical assistance. Special thanks to Ono Pharmaceutical Co, Ltd. for the provision of sivelestat as the study medication.

## AUTHOR CONTRIBUTIONS

All authors designed research. H.M. and S.-O. S. performed research. T.Y., N.H., E. H., C.-I.H., M.-A.O., K. S., T. Y., and T.N. provided assistance. H.M. and S.-O. S. analyzed data. H.M. and T. Y. wrote the paper. All authors approved the final manuscript.

